# VEGFC induced cell cycle arrest mediates sprouting and differentiation of venous and lymphatic endothelial cells

**DOI:** 10.1101/2020.06.17.155028

**Authors:** Ayelet Jerafi-Vider, Noga Moshe, Gideon Hen, Daniel Splittstoesser, Masahiro Shin, Nathan Lawson, Karina Yaniv

**Author notes:** Corresponding Author, Karina Yaniv, Department of Biological Regulation, Weizmann Institute of Science, Rehovot, 76100, Israel.

## Abstract

The emergence and growth of new vessels requires a tight synchronization between proliferation, differentiation and sprouting, traditionally thought to be controlled by mitogenic signals, especially of the VEGF family. However, how these cues are differentially transduced, by sometimes even neighboring endothelial cells (ECs), remains unclear. Here we identify cell-cycle progression as a new regulator of EC sprouting and differentiation. Using transgenic zebrafish illuminating cell-cycle stages, we show that venous and lymphatic precursors sprout from the Cardinal Vein (CV) exclusively in G_0_/G_1_, and reveal that cell-cycle arrest is induced in these ECs by p53 and the CDK-inhibitors p27 and p21. Moreover, we demonstrate that *in vivo*, chemical and genetic cell-cycle inhibition, results in massive vascular growth. Mechanistically, we identify the mitogenic VEGFC/VEGFR3/ERK axis as direct inducer of cell-cycle arrest in angiogenic ECs and characterize the cascade of events governing venous vs. lymphatic segregation and sprouting. Overall, our results uncover an unexpected mechanism whereby mitogen-controlled cell-cycle arrest boosts sprouting, raising important questions about the use of cell-cycle inhibitors in pathological angiogenesis and lymphangiogenesis.

## Introduction

The formation of a new vessel from a pre-existing one requires a tightly regulated synchronization between different processes, such as cell proliferation, specification and motility^1,2^. The balance between cell proliferation and differentiation is considered a hallmark of cell fate determination during embryonic development ^3,4^. In particular, the G1 phase of the cell cycle acts as a critical checkpoint in cell fate decisions, including stem cell differentiation^5,6^. The length of the G1 stage is known to be controlled by cell cycle inhibitors such as p21 and p27, which suppress cyclin\CDK complex activity^7–9^. While the link between cell proliferation and differentiation has been extensively investigated in various embryonic tissues^4^, it is only recently that it attracted attention in the context of vascular formation. In the mouse retina for instance, shear stress-dependent upregulation of Notch signaling was shown to induce elevation of p27 and blockage of cell cycle in the G1 phase, which in turn lead to arterial specification^10^. Cell proliferation is also tightly linked to angiogenic sprouting. Both in development^11,12^ and pathologies^13,14^, VEGF stimulates both sprouting and growth of endothelial cells (ECs) and formation of new blood vessels, acting mainly through its receptor VEGFR2. Surprisingly however, a recent report demonstrated that high levels of VEGF-A in fact inhibit the proliferation of arterial tip cells in the postnatal mouse retina, while promoting their active migration and sprouting^15^. In contrast to these advances, little is known about the spatiotemporal dynamics of cell cycle progression in ECs *in vivo* and how it affects angiogenesis and sprouting. Moreover, whether cell cycle progression is linked to the initial venous (VEC) vs. lymphatic (LEC) segregation has not been addressed.

During embryonic development, VECs and LECs sprout from the cardinal vein (CV) towards a gradient of VEGFC^16,17^. Accordingly, VEGFC-deficient mice and zebrafish fail to establish a proper lymphatic system^16,18–22^. VEGFR3 (also known as Flt4) is the main receptor for VEGFC, and its activation mediates LEC proliferation, migration and survival^23–25^. In zebrafish mutations in the *vegfr3* gene result in complete absence of the lymphatic vasculature, without affecting blood vessel sprouting^22,26,27^. In addition to its mitogenic role, VEGFC has also been shown to control LEC differentiation, through ERK activation^28,29^. ERK phosphorylation, which is commonly known to transduce mitogenic signals, is augmented downstream to VEGFC-FLT4 signaling to promote both differentiation and sprouting of lymphatic progenitors^28,29^. Nevertheless, how the processes of EC proliferation, cell fate decision (e.g. venous vs. lymphatic differentiation), and sprouting are orchestrated during embryonic development to enable two different vessel types to arise almost simultaneously from the same vessel-the PCV-, remains largely unknown.

Here we investigate the link between cell cycle progression, differentiation and sprouting of VECs and LECs emerging from the PCV of the zebrafish embryo. Using live imaging of transgenic reporters highlighting different stages of the cell cycle we show that both VECs and LECs bud from the dorsal side of the PCV in G_0_/G_1_ phase, and demonstrate that cell cycle arrest is specifically induced in these “angiogenic” ECs through upregulation of p53, p21 and p27. Using chemicals and genetic manipulations we further reveal that induction of cell cycle arrest in PCV ECs results in massive exit of undifferentiated ECs from the PCV and excessive sprouting. Molecularly, we identify the Vegfc-Flt4-ERK signaling axis as upstream regulator of p53, p21 and p27 expression in sprouting ECs, and characterize the full cascade of events orchestrating the segregation of venous and lymphatic vessels. Overall, our results uncover a new layer of regulation of sprouting angiogenesis whereby inhibition of cell cycle progression activates sprouting, raising important questions about the role of cell cycle inhibitors in states of pathological angiogenesis.

## Results

### Endothelial cells sprout from the PCV in G_0_/G_1_ phase

In order to investigate the link between cell cycle dynamics and EC sprouting *in vivo*, we established a *Tg(fli1:gal4;uasKaede;uas:fucci)- “fli:fucci”* reporter (Figure 1a) in which EC nuclei in G_0_/G_1_ phase are labeled in red, and EC nuclei in S-M phases display green fluorescence^30^. Confocal imaging of transgenic embryos revealed that neither the total number of PCV cells (Figure S1a), nor the relative number of G_0_/G_1_ red cells in the PCV (Figure 1b-f) changed between 26-42 hpf. In contrast, we detected significant differences in the location of G_0_/G_1_ ECs between these two time points. While before lympho-venous sprouting (~26 hpf) G_0_/G_1_ cells were evenly distributed throughout the PCV (Figure 1b,c,g), during active sprouting (~40 hpf), ~80% of the G_0_/G_1_ population was detected in its dorsal side (Figure 1d,e,g). We used long-term time-lapse imaging of *fli:fucci* embryos to track the dynamics of cell cycle progression during PCV sprouting (Fig. 1h-o, Figure S1b and Supplementary Movie 1). We observed that ECs bud from the PCV in G_0_/G_1_ phase (red) and upon crossing the anatomical level of the dorsal Aorta (DA) they turn green (Figure 1k,n,o, white arrowheads), indicating entry into S-M phases of the cell cycle. This process was independent of whether the sprouting cell was a VEC that gave rise to a venous intersegmental vessel (vISV) (Figure 1h-k, Figure S1b,c), or a lymphatic progenitor that incorporated into the parachordal chain (PAC) (Figure 1l-o, Figure S1b,c), highlighting a significant correlation between cell cycle progression and global PCV sprouting.

**Figure 1.**
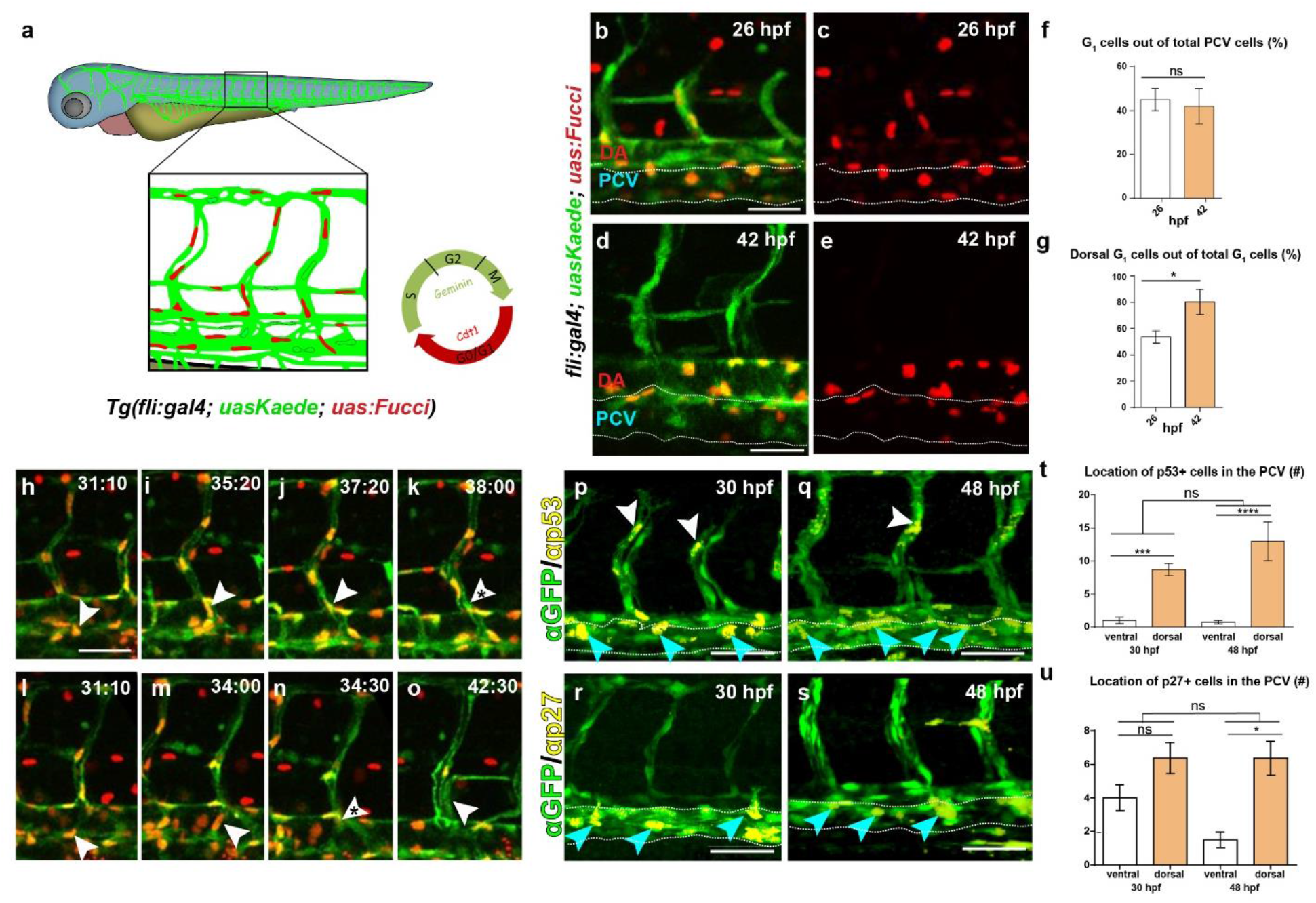
Endothelial cells sprout from the PCV in G_0_/G_1_ phase. (**a**) Schematic representation of cell cycle stages in ECs as highlighted by the *Tg(fli1:gal4;uas:kaede;uas:fucci)* reporter. (**b-e**) Selected confocal snapshots from a time-lapse series of a *Tg(fli1:gal4;uas:kaede;uas:fucci)* embryo depicting the distribution of G_0_/G_1_ ECs in the PCV (outlined by dashed lines) at 26 (**b,c**) and 42 (**d,e**) hpf. (**f**) Fraction of G_0_/G_1_ ECs out of total PCV cells (n=4). (**g**) Spatial distribution of G_0_/G_1_ ECs in the PCV at 26 and 42 hpf (n=4). (**h-o**) Selected confocal snapshots from a time-lapse series of a *Tg(fli1:gal4;uasKaede;uas:fucci)* embryo showing two different ECs leaving the PCV in G_0_/G_1_ stage (red nuclei, white arrowheads) and re-entering cell cycle (**k,n**; asterisks). (**p-s**) Confocal images of 30 (**p,r**) and 48 (**q,s**) hpf *Tg(fli1:EGFP)* embryos immuno-stained for p53 and anti-GFP (p,q; blue arrowheads point to double stained ECs in the PCV) or for p27 and anti-GFP (r,s; blue arrowheads point to double stained ECs in the PCV). (**t**) Location of p53+ (n30 hpf=7, n48 hpf=4) and (**u**) p27+ (n30 hpf=18, n48 hpf=8) ECs in the PCV (outlined by dashed lines). Scale bars:,**b-e,h-o=**40μm; **p-s**=70μm; *p<0.05, ***p<0.001, ****p<0.0001, ns; not significant.

### G_0_/G_1_ cell cycle arrest in PCV ECs is mediated by p53, p21 and p27

Cell cycle progression is coordinated through a complex molecular network wherein each phase is controlled by specific Cyclin-dependent kinases (CDKs) and their binding Cyclins. While Cyclins activate CDKs, Cyclin-CDK inhibitors (CKIs), such as p16, p15, p27 and p21, negatively regulate CDK activity and cell cycle progression^9^, inducing growth arrest at the G_0_/G_1_ checkpoint.

Similarly, p53 activation leads to G_0_/G_1_ cell cycle arrest associated with apoptosis, cell differentiation^31,32^ and migration^33^.

In order to investigate the molecular mechanisms inducing cell cycle arrest in sprouting ECs we analyzed the expression of p53, p27 and p21 in 30 and 48 hpf *Tg(fli1:EGFP)* embryos. Interestingly, antibody staining revealed clear accumulation of p53 in “angiogenic” ECs located in the dorsal PCV (Figure 1p,q,t, blue arrowheads), as well as in arterial tip cells (Figure 1p,q, white arrowheads). Similar results were obtained following analysis of p27 expression (Figure 1r,s,u, blue arrowheads) and *in situ* hybridization for *p21* (data not shown).

The above results prompted us to investigate the effects of forcing cell cycle arrest, on PCV sprouting. To this end, we treated 20 hpf *fli:fucci* embryos with Roscovitine, a well-established p53 stabilizer and CDK inhibitor^34–36^. As expected, Roscovitine treatment resulted in increased numbers of G_0_/G_1_ mCherry+ cells, that were evenly distributed in the PCV of treated embryos by 48 hpf (Figure 2a-c). Surprisingly however, PCV sprouting was not reduced, but rather enhanced in these embryos, which displayed abnormal and ectopic sprouts and exhibited excessive branching as compared to DMSO-treated siblings (Figure 2d-f). To further confirm these results, we evaluated the ability of additional cell cycle inhibitors to recapitulate these phenotypes. 20 hpf *Tg(lyve1b:dsRed2;fli1:EGFP)* embryos were treated with either Aphidicoline (S), Nocodazole (M), Etoposide (G_2_), or Flavopiridol (G_0_/G_1_) as previously described^37^, and phenotypes were assessed at 3 dpf (Figure 2g-j). As expected, both Aphidicoline and Nocodazole treatments affected global embryonic development and therefore resulted in an unhealthy vasculature and severe sprouting defects (Figure 2g,h), whereas Etoposide, which inhibits cell cycle at the G_2_ phase, did not affect PCV sprouting whatsoever (Figure 2i). In contrast to these phenotypes, Flavopiridol treatment (G_0_/G_1_ inhibitor) induced the formation of ectopic PCV sprouts (Figure 2j, white arrowheads), resembling those observed following Roscovitine supply (Figure 2e).

**Figure 2:**
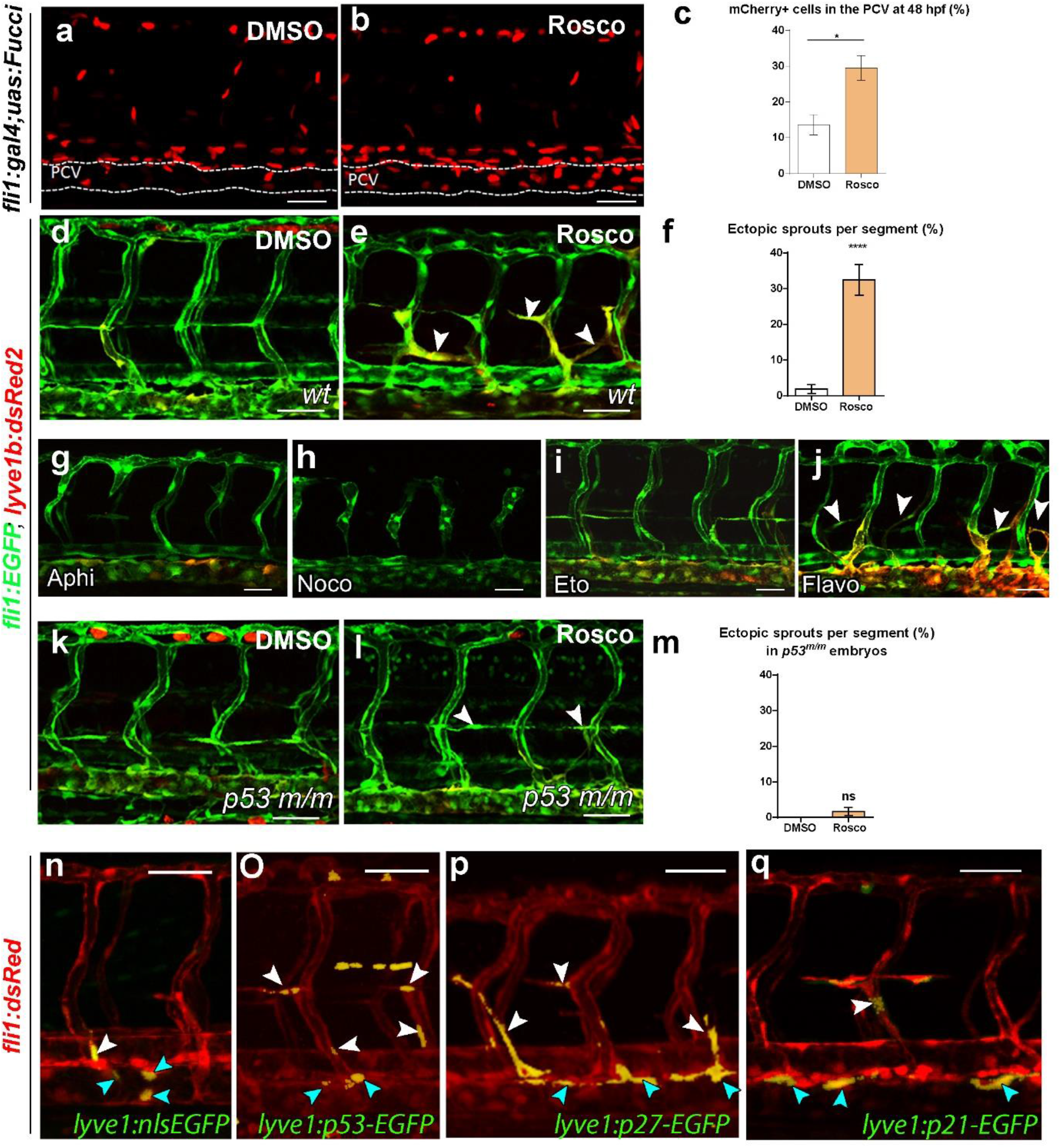
G_0_/G_1_ cell cycle arrest in PCV ECs is mediated by p53-p21 and p27. (**a,b**) Confocal images of *Tg(fli1:gal4;uas:fucci)* (red channel) embryos at 48 hpf, showing increased numbers of G_0_/G_1_ *mCherry+* nuclei in the PCV (outlined by dashed lines) following Roscovitine vs. DMSO treatment; quantified in **c** (n_DMSO_=12, n_Rosco_=10). (**d,e**) Confocal images of *Tg(fli1:EGFP;lyve1b:dsRed2)* embryos showing ectopic and mis-patterned PCV sprouts at 48 hpf (**e**, arrowheads), that are not detected in DMSO-treated siblings (**d**). (**f**) Number of ectopic sprouts per segment in Rosco vs. DMSO-treated embryos (n_DMSO_=12, n_Rosco_=12). (**g-j**) Confocal images of 3 dfp *Tg(fli1:EGFP;lyve1b:dsRed2)* embryos treated with Aphidicolin (**g**, n=6), Nocozadole (**h**, n=11), Etoposide (**i**, n=11) and Flavopiridol (**j**, n=14), showing ectopic and mis-patterned PCV sprouts following Flavopiridol treatment (**j**, arrowheads). (**k,l**) Confocal images of *Tg(fli1:EGFP;lyve1b:dsRed2;p53*^m/m^) embryos at 48 hpf, showing normal PACs following DMSO (**k**) and Roscovitine (**l,** arrowheads) treatments. (**m**) Percentage of ectopic sprouts per segment in Roscovitine (n=12) vs. DMSO (n=12) treated *p53^m/m^* embryos. (**n-p**) Confocal images of *Tg(fli1:DsRed)* embryos 48 after injection with *lyve1:nEGFP* (**n**, n=12), *lyve1:p53-EGFP* (**o**, n=30) or *lyve1:p27-EGFP* (**p**, n=12). (**q**) Confocal images of *Tg(fli1:DsRed;lyve1:p21-EGFP)* embryos at 48 hpf. Blue arrowheads in **n-q** point to GFP+ ECs in the dorsal PCV, white arrowheads denote GFP+ lympho-venous sprouts. Scale bars: **a-l**=40 μm, **n-q**=50 μm. *p<0.05, ****p<0.0001, ns; not significant.

Since Roscovitine is known to halt progression through the cell cycle by selective inhibition of CDKs and by stabilization of p53^34–36^ we assessed its ability to induce ectopic sprouting in *p53* mutant embryos^38^. Unlike WT animals, Roscovitine-treated *p53* mutants did not display any noticeable defects (Figure 2k-m), strongly suggesting that the excessive sprouting phenotype is p53 dependent. Of note, *p53* mutant embryos develop normally, and possess normal trunk vasculature (Supplementary Figure 2a,b), hinting at the presence of alternative compensatory mechanisms. We next set out to investigate whether G_0_/G_1_ arrest can cell-autonomously drive EC sprouting from the PCV. To this end, we over-expressed EGFP-tagged forms of p53, p27 or p21 in the PCV of *Tg(fli1:DsRed)* embryos using the *lyve1* promoter, and analyzed the behavior of the labeled cells at 48 hpf. A *lyve1:nlsEGFP* plasmid was used as control. As seen in Figure 2n-q, all injected plasmids were specifically expressed as expected, in the PCV and in venous and lymphatic sprouts at 48 hpf. Interestingly however, assessment of the location of the GFP+ cells at 48 hpf, revealed significant differences between the injected groups. While only 25.66% of EGFP+ cells were detected outside the PCV (i.e vISVs and PACs) in control-injected embryos (Figure 2n and Figure S2c), almost twice the amount (41.92% and 43.87%) was observed in venous and lymphatic sprouts following overexpression of p53 and p27, respectively (Figure 2o,p white arrows and Figure S2c). In addition, there was a significant shift of GFP+ cells from the ventral to the dorsal side of the PCV, suggesting that p53 and p27 act cell-autonomously in ECs to induce sprouting. In contrast to p53 and p27 overexpressing embryos, *lyve1:p21-EGFP* injected *larvae* survived through adulthood and were fertile, enabling generation of stable transgenic fish and assessment of the spatial location of EGFP+ cells in their progeny at 48hpf. Notably, while only 4.65% of the EGFP+ cells in p21 overexpressing embryos were located in the ventral PCV (Figure 2q and Figure S2c), the vast majority was detected in the dorsal PCV (67.44%) and in lymphatic/venous sprouts (27.9%). Taken together these results indicate that increased expression of p53, p27 and p21 acts cell-autonomously in PCV ECs to promote cell cycle arrest-induced sprouting.

### Cell cycle arrest in dorsal PCV ECs is VegfC/VegfR3-dependent

During embryonic development, venous and lymphatic ECs sprout from the cardinal vein (CV) in response to Vegfc-Vegfr3 signaling^17,39^. In zebrafish, Vegfc is expressed in the hypochord and the Dorsal Aorta (DA)^40–42^ and is essential for sprouting of both lymphatic and venous ECs from the PCV^18^. Vegfr3/Flt4, the main receptor for VegfC, is expressed by blood and lymphatic ECs, both in mouse^24^ and zebrafish^42–44^, and is required for remodeling of the vascular network.

We undertook several approaches in order to define the epistatic relationship between cell cycle arrest-induced sprouting and the Vegfc/Flt4 axis. First, we examined whether Roscovitine treatment affects the expression of these two factors. *In situ* hybridization at 26 hpf showed no differences in *flt4* and *vegfc* mRNA levels between Roscovitine vs. DMSO treated embryos (Figure S3a-d). Next, we asked whether cell cycle arrest is sufficient to induce ectopic sprouting in the absence of active Vegfc-Flt4 signaling. Interestingly, addition of Roscovitine to the water of *vegfc*^18^ and *flt4^28^* mutants, which completely lack PCV-derived sprouts, did not revert this phenotype (Figure 3a-g), suggesting that Vegfc-Flt4 signaling is required in order to enable cell cycle arrest-induced sprouting. Lastly, we asked whether, in spite of being Vegfc a major pro-proliferative and survival factor, it can in fact trigger cell cycle arrest in ECs, to promote sprouting. Indeed, immunostaining revealed a drastic reduction in the number of p27+ and p53+ ECs in the PCV of both *vegfc* and *flt4* mutant embryos as compared to wt sibling (Figure 3h-o, blue arrows and Figure S3e-j), indicating that the specific expression of these CKIs in angiogenic ECs is Vegfc-Flt4-dependent.

**Figure 3:**
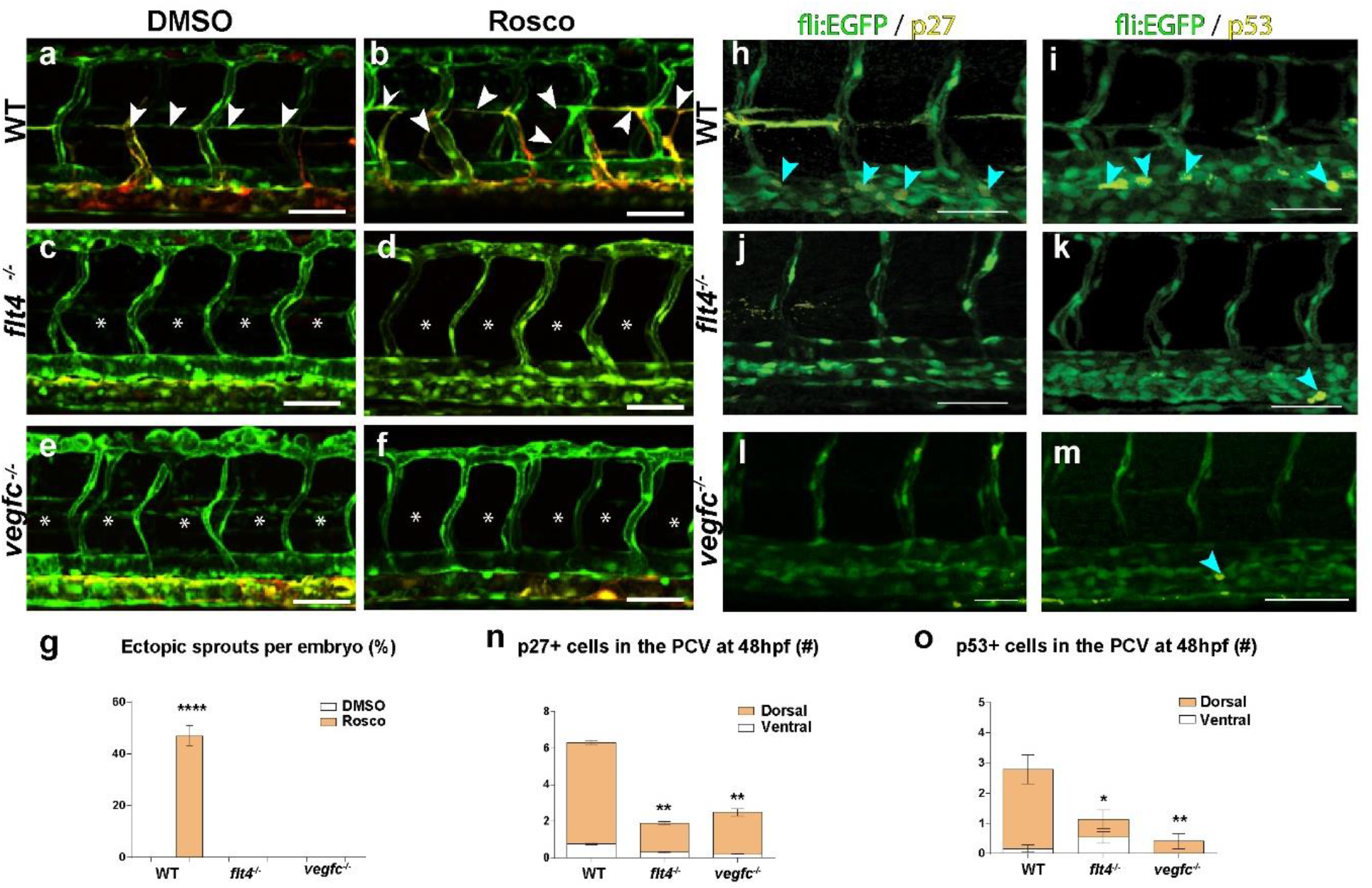
VegfC/VegfR3 induce cell cycle arrest in dorsal PCV ECs. (**a-f**) Confocal images of WT (**a,b**), *flt4^-/-^* (**c,d**) and *vegfc^-/-^* (**e,f**); *Tg(fli1:EGFP;lyve1b:dsRed2)* embryos at 3 dpf, treated with DMSO (n_WT_=24, n_*flt4-/-*_ =10, n_*vegfc-/-*_= 6) or Roscovitine (n_WT_=19, n_*flt4-/-*_ =8, n_*vegfc-/-*_=10). White arrowheads in **a,b** point to normal and ectopic lympho-venous sprouts that are absent in *flt4* and *vegfc* mutants (**c-f**, asterisks); quantified in (**g**). (**h-m**) Confocal images of WT, *flt4^-/-^* and *vegfc^-/-^; Tg(fli1:EGFP)* embryos at 48 hpf immunostained with p27 (**h,j,l**) or p53 (**i,k,m**) antibodies. Pictures show co-localization channel in yellow; blue arrowheads point to immunostained ECs. (**n,o**) Number of p27 (**n**) and p53 (**o**) stained ECs in the dorsal vs. ventral PCV of WT, *flt4^-/-^* and *vegfc^-/-^; Tg(fli1:EGFP)* embryos (n_p27 wt_=19, n_p27 *ttt4-/-*_=8, n_p27 *vegfc*-/-_=10, n_p53 WT_=17, n_p53 *flt4-/-*_=15, n_p53 *vegfc*-/-_=5). *p<0.05, ***p<0.001, ****p<0.0001, ns; not significant. Scale bars=70 μm.

Recently it has been shown that phosphorylated ERK (pERK) is detected in the PCV at the onset of sprouting, and that ERK phosphorylation is required for both venous-lymphatic sprouting and lymphatic differentiation, downstream of Vegfc/Flt4. Accordingly, blocking ERK phosphorylation and activation, by chemical inhibition of MAPK/ERK Kinase (MEK), abolished PCV sprouting^28^. Based on these and our results, we wondered how two opposing outputs such as cell cycle arrest on the one hand, and ERK activation (typically a mitogenic signal) on the other, can be achieved in sprouting ECs downstream of Vegfc/Flt4 signaling. To answer this question, we utilized a well-known selective inhibitor of MEK1/2, - SL327 - which blocks ERK phosphorylation thus inhibiting pERK-dependent signaling^28^. Treatment of zebrafish embryos with SL327 at 24 hpf was shown to inhibit budding of both venous and lymphatic sprouts^28^. We treated 20 hpf *fli:fucci* and *Tg(mrc1a:EGFP)* ^45^ embryos with either Roscovitine or DMSO, then added SL327 at 24 hpf, and analyzed PCV sprouting one day later. As predicted, both pharmacological inhibitors suppressed cell proliferation as confirmed by the increased number of red nuclei detected in *fli:fucci* treated embryos as compared to their DMSO-treated siblings (Figure S4a-d). Likewise, both treatments yielded the expected phenotypes; namely, Roscovitine induced excessive PCV sprouting (Figure 4a,b,e) and SL327 abolished almost completely the emergence of venous and lymphatic sprouts from the PCV (Figure 4a,c,e). To our surprise however, the combination of Roscovitine and SL327 treatments, resulted in a remarkable rescue of PCV sprouting (Figure 4d,e), suggesting that G_0_/G_1_ cell cycle arrest acts downstream to ERK signaling to enable PCV sprouting. Likewise, stable overexpression of p21 in *Tg(mrc1a:EGFP;lyve1:p21-EGFP)* embryos (Figure 4f-j), was sufficient to restore PCV sprouting in the absence of active MEK signaling as well. Finally, analysis of p27 (Figure 4k-m, Figure S4e-f) and p53 (Figure 4n-p, Figure S2g-h) expression following SL327 treatment revealed a significant decrease in the numbers of PCV-ECs positively stained for these cell cycle inhibitors, uncovering a novel mechanism whereby pERK controls PCV sprouting, through induction of cell cycle arrest.

**Figure 4:**
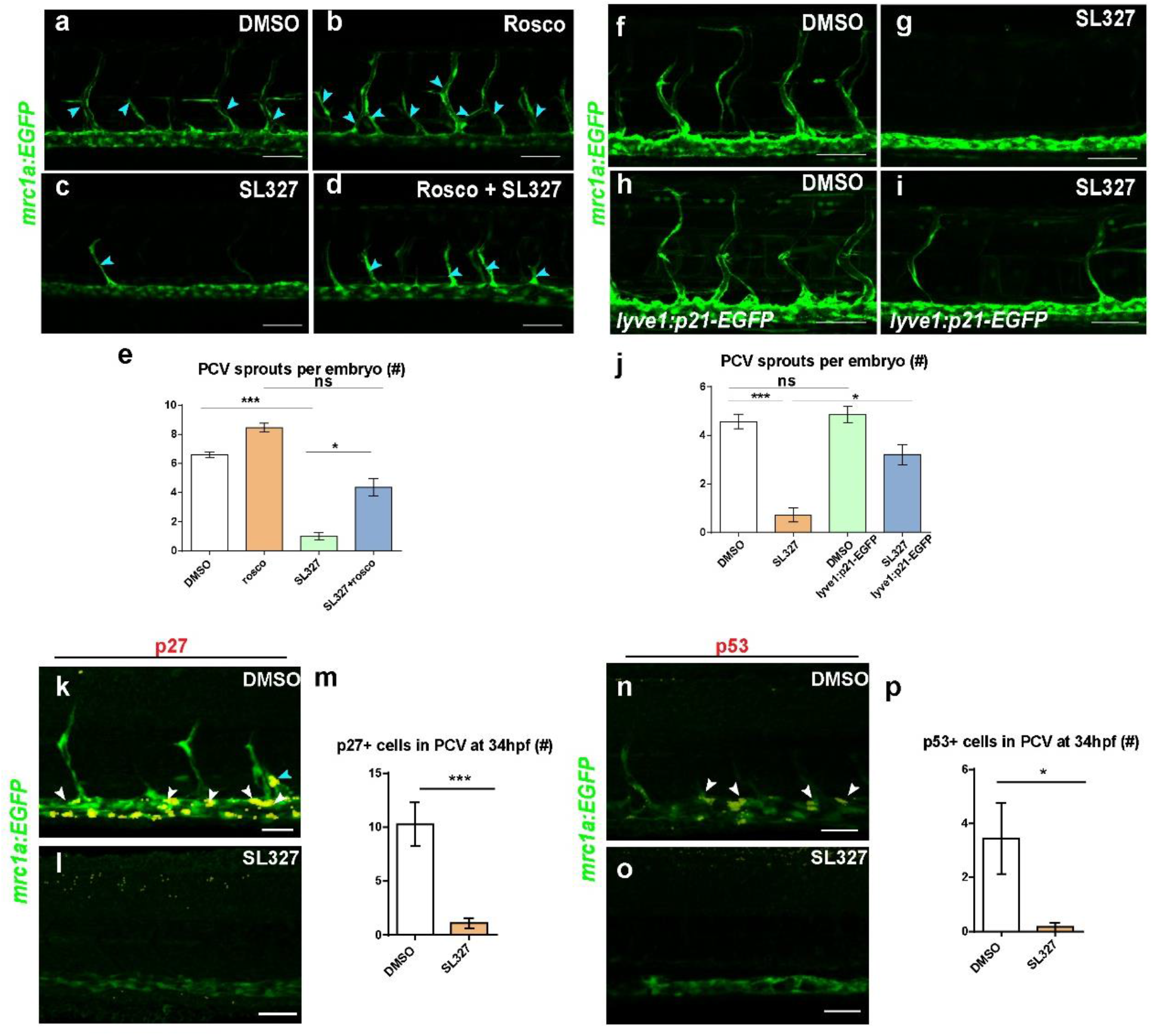
Cell cycle arrest in dorsal PCV cells is pERK dependent. (**a-d**) Confocal images of *Tg(mrc1a:EGFP)* embryos at 48 hpf treated with DMSO (**a**), Roscovitine (**b**), SL327 (**c**) or Roscovotine+SL327 (**d**) showing partial rescue of lympho-venous sprouting following Roscovotine+SL327 (**d**) as compared to SL327 only treatment (**c**), quantified in **e** (n_DMSO_=12, n_Rosco_=13, n_SL327_=17, n_Rosco_+SL327=16). Blue arrowheads in **a-d** point to PCV emerging sprouts. (**f-j**) Confocal images of *Tg(mrc1a:EGFP)* (**f,g**) and *Tg(lyve1:p21-EGFP;mrc1a:EGFP)* (**h,i**) embryos treated with DMSO (**f,h**) or SL327 (**g,i**) showing partially restored lympho-venous sprouting in *lyve1:p21-EGFP* overexpressing embryos (**i**), quantified in **j** (n_DMSO_=7, n_SL327_= 7, n_*lyve1:p21-EGFP*+DMSO_=7, n_*lyve1:p21-EGFP*+SL327_=15). (**k-p**) Confocal images of *Tg(mrc1a:EGFP)* embryos at 34 hpf, showing reduced expression of p27 (**k,l**) and p53 (**n, o**) following SL327 treatment, quantified in **m,p** (n_DMSO p27_=7, n_SL327 p27_=8, n_DMSO p53_=9, n_SL327 p53_=6). Co-localization channel is shown in yellow. *p<0.05, ***p<0.001. Scale bars=30μm

### Forced G_0_/G_1_ cell cycle arrest affects venous-lymphatic differentiation

Given that the VegfC-VegfR3-ERK cascade was found to control not only venous and lymphatic sprouting, but also lymphatic specification^28,29,46^, we decided to investigate whether cell cycle progression plays also a role in this process. We began by analyzing the expression of venous and lymphatic markers following Roscovitine treatment (Figure 5). Indeed, *in situ* hybridization revealed complete absence of *lyve1* and *nr2f2* mRNA expression in treated embryos (Figure 5a,b). Moreover, the number of *prox1+* cells in the PCV of Roscovitine-treated *Tg(fli1:EGFP;prox1a:KalTA4-4xUAS-E1b:uncTagRFP)^47^* embryos was significantly reduced as compared to DMSO-treated siblings (Figure 5c-e and Figure S5e,f). These results suggest that forced cell cycle arrest inhibits the segregation between venous and lymphatic progenitors within the PCV, while promoting their sprouting and migration in an undifferentiated state. To better understand this phenotype, we time-lapse imaged *Tg(fli1:EGFP)* embryos continuously exposed to Roscovitine, and tracked the behavior of ECs sprouting from the PCV. As seen in Figure 5f-k (also in Supplemental Movie 2), we detected single sprouts where the leading cell (colored in red), connects to a neighboring arterial ISV to generate a lumenized vISV (Figure 5h,I, Supplemental Movie 2), whereas the following cell (colored in blue), divides horizontally and generates two daughter cells, one of them incorporating into the PAC, while the other joins a vISV (Figure 5i-k, Supplemental Movie 2). This phenotype of PCV cells generating both venous and lymphatic cells after sprouting, is not normally observed in WT embryos, as cells acquire their lymphatic fate prior to leaving the PCV^47,48^.

**Figure 5:**
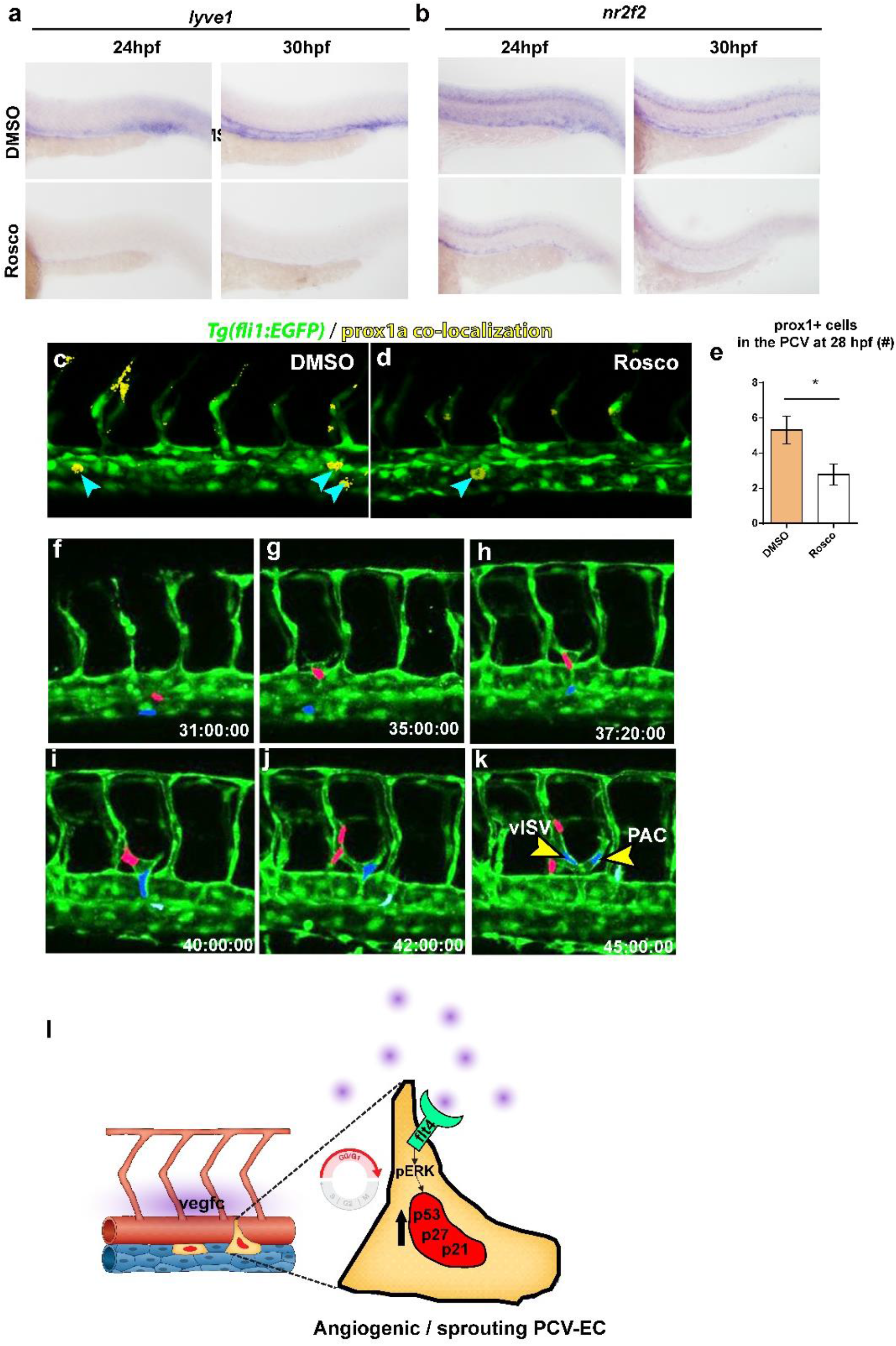
Forced G_0_/G_1_ cell cycle arrest affects venous-lymphatic differentiation. (**a,b**) *In situ* hybridization at 24 and 30 hpf showing reduced expression of *lyve1* (**a**) and *nr2f2* (**b**) following Roscovitine treatment. (**c,d**) Confocal images of 28 hpf *Tg(fli1:EGFP;prox1a:kalt4:UAS:uncTagRFP)* embryos showing reduced numbers of *prox1+* cells (blue arrowheads) within the PCV upon Roscovitine treatment. Co-localization channel is shown in yellow, quantified in **e**. (**f-k**) selected images from a time-lapse series depicting the dynamics of PCV sprouting in the presence of Roscovitine. 2 ECs (colored in red and blue) leave the PCV as part of a single sprout (**h**) that generates two vessel types. The leading EC (red) connects to an arterial ISV to generate a vISV (**i,j**), while the following cell (blue) divides (**j**) and gives rise to 2 daughter cells (**k**, arrowheads) one joins a vISV and the second one incorporates into the nascent PAC. (**l**) Schematic model depicting cellular events that lead to cell cycle arrest induced sprouting. Dorsal cells of the PCV sense high levels of Vegfc (purple) secreted from the hypocord and the DA. Activation of vegfr3/flt4-ERK signaling in these cells leads to cell cycle arrest-induced sprouting.

Taken together, our results illustrate how different signaling pathways are exquisitely synchronized both in time and space to support proper angiogenesis and lymphangiogenesis in the developing embryo and introduce a novel level of regulation to this process involving cell cycle arrest (Figure 5l). Based on our findings we propose that distinct levels of VegfC/ERK control both sprouting and differentiation of PCV-ECs, through differential regulation of cell cycle progression. Differentiated venous and lymphatic ECs located in the dorsal side of the PCV, are exposed to high levels of the VegfC ligand, which is secreted from the hypochord and the DA. In these cells, ERK signaling (most likely of high magnitude and prolonged activation) induces the expression of p53, p21 and p27, promoting sprouting. Concordantly, the expression of the cell cycle inhibitors was drastically reduced in the absence of ERK and both Roscovitine treatment and cell-autonomous overexpression of p21 were sufficient to overcome the inhibition of sprouting following SL237 treatment. As the sprouting cells migrate dorsally and cross the anatomical level of the DA they lose p53, p21 and p27 expression, which triggers cell cycle re-entry. In contrast, cells in the ventral side of the PCV, which we have previously show are mostly undifferentiated angioblasts^47^, perceive low levels of VegfC and lesser ERK activation, impeding CDK inhibitors’ expression, thereby enabling angioblast proliferation and LEC differentiation.

## Discussion

Here we show that cell cycle progression plays a crucial role in sprouting and differentiation of venous and lymphatic ECs. Illuminating cell cycle progression in ECs, through the use of *Tg(fli1:gal4;uasKaede;uas:fucci)* embryos, provided valuable insight into the normal dynamics of cell proliferation in the developing vasculature. We found that during active sprouting (30–42 hpf) G_0_/G_1_ cells populate the dorsal side of the PCV. These findings correlate with the accumulation of p53, p27 and p21 expressing cells in the dorsal PCV at sites of sprouting, reinforcing the idea that cell cycle arrest is required in these cells in order to enable sprouting.

We also show that arresting cell cycle in G_0_/G_1_ by using Roscovitine, a well-established Mdm2 and CDK inhibitor, or Flavopiridol, a selective potent CKD inhibitor clinically efficient for cancer treatment^49,50^, induces the formation of undifferentiated PCV sprouts. These results support the idea that cell cycle regulation, particularly the regulation of the G_0_/G_1_ phase, plays essential roles in proper sprouting and differentiation of LECs. Interestingly, these chemical treatments were unable to induce excessive sprouting in *p53* mutants, suggesting the specific involvement of p53 in this process. Using genetic manipulations, we demonstrate that PCV ECs overexpressing p53, p27 and p21 acquire an “angiogenic” phenotype and become more prone to sprout, indicating a cell-autonomous requirement of cell cycle arrest to enable sprouting.

During embryonic development, venous and lymphatic ECs sprout from the PCV in response to Vegfc-Vegfr3-ERK signaling^17^. Our results indicate that this axis in fact induces cell cycle arrest in dorsal PCV ECs, to promote their sprouting and migration. Both *vegfc* and *flt4* mutants display reduced number of p53 and p27 expressing cells within the PCV. Accordingly, we were unable to induce sprouting in these mutants following treatments with CDIs, suggesting that Vegfc and its Flt4 receptor are responsible for cell cycle arrest induction in sprouting ECs. Downstream to Vegfc-Vegfr3, ERK was shown to induce both differentiation and sprouting of lymphatic vessels^28,29^. While ERK phosphorylation plays a pivotal role in diverse cellular functions, including cell proliferation, differentiation, migration and survival^51,52^, ERK activation can also elicit opposite outcomes, such as cell-cycle arrest and cell death depending on the context^53^. Recent progress has demonstrated that differences in the duration and magnitude of ERK activity generate variations in signaling output that regulate cell fate decisions^53,54^. In the case of ECs, ERK phosphorylation is mostly associated with proliferation and sprouting^28,55,56^. Our results in contrast, suggest that ERK phosphorylation drives sprouting and migration of PCV ECs by inducing cell cycle arrest. Accordingly, both Roscovitine treatment and EC-specific overexpression of p21 reverted the inhibition of sprouting exerted by the potent ERK phosphorylation inhibitor SL327. These results are in line with recent findings demonstrating that high VEGF signal induces cell cycle arrest in sprouting tip cells of the mouse retina, through pERK dependent elevation of p21^15^.

Overall, our results uncover a new layer of regulation of venous vs. lymphatic EC sprouting and differentiation, whereby a mitogenic signal induces cell cycle arrest to enable angiogenesis. These findings have important implications for the putative effects of using cell cycle inhibitors in settings of pathological angiogenesis, including cancer.

## Materials and methods

### Zebrafish husbandry and transgenic lines

Zebrafish were raised by standard methods and handled according to the guidelines of the Weizmann Institute Animal Care and Use Committee^57^. For all imaging, *in situ* hybridization and immunofluorescence staining procedures, embryos were treated with 0.003% phenylthiourea (PTU, Sigma-Aldrich) from 8 hpf to inhibit pigment formation. *Tg(fli1:EGFP)^yl^* ^58^, *Tg(lyve1b:dsRed2)^nz101^* ^59^, *Tg(fli1:dsRed)^um13^* ^60^, *Tg(fli1:gal4^ubs3^;uasKaede^rk8^)^47^, tp53^m214k/m214k^* ^38^, *Tg(mrc1a:EGFP)^y251^* ^45^ *TgBAC(prox1a:KalT4-UAS:uncTagRFP)^nim^* ^47^, *vegfc^um18/ um18^* ^18^ and *flt4^um203/um203^* ^27^ were previously described.

To generate the “*fli:fucci*” transgenic reporter the ubiquitous promoter in the dual *fucci* construct^30^ was replaced with the UAS sequence. The *uas:cerulian-zGeminin-2A-Cherry-zCdt1Pa* construct was then injected into *Tg(fli1:gal4^ubs3^;uasKaede^rk8^)* embryos at 1-cell stage along with Tol2 transposase mRNA.

We used the following primers to amplify the full-length coding sequences of zebrafish *p53, p27* and *p21*: *p53*- 5’-TTCACAGCAATGGCGCAAAACG-3’ and 5’-ACATAGCGTCAAGGGGTTTACTGG-3’, *p27*- 5’-ATGTCCAATGTTCGCTTGTCT-3’ and *5’-CATTGAGTCCGACACCCACA-3’* 3’- TGTGGGTGTCGGACTCAATG-5’, *p21*- 5’-CTGATACTGCTCCTGAGGAGATCTGA-3’ and5’- CTACGAGACGAATGCAGCTCCAGACA -3’. After cloning and sequencing, a Gateway-compatible (Invitrogen) middle entry clone was generated using Gateway BP clonase mediated recombination. The *p53, p27* and *p21* coding sequences were then transferred into a *pDestTol2pA2* vector^61^ along with a *p5E-lyve1* fragment^59^, using Gateway LR clonase (Invitrogen) mediated reaction. The *lyve1:nEGFPpA* construct was generated by combining the *p5E-lyve1* promoter with nuclear localization signal (nls)-EGFP middle entry (*pME-nEGFP*) and *p3E-pA* constructs. The final *lyve1:nEGFPpA*, *lyve1:p53-EGFPpA, lyve1:p27-EGFPpA and lyve1:p21-EGFPpA* constructs (30pg) were co-injected with *Tol2* transposase mRNA (30pg) into 1-cell stage *Tg(fli1:DsRed)^um13^* embryos.

### Chemical treatments

The following chemicals were dissolved in DMSO and added to the fish water at 20 hpf: Roscovitine (Santa Cruz Biotechnology, cat# sc-24002A), 50μM; Flavopiridole^37^ (Enzo life sciences, ALX-430-161-M005), 5nM; Etoposide^37^ (Sigma Aldrich, E1383), 1000μM; Nocozadole^37^ (Sigma, M1404), 300nM; Aphidicoline^32^ (Sigma, A0781), 297μM; SL327 (Sigma, S4069), 15 μM^28^. Controls were incubated with 0.1% DMSO.

### *In situ* hybridization and immunostaining

*In situ* hybridization was performed as described^47^. The following primers were used to generate the corresponding riboprobes:

*flt4:* 5’- TGGAGTTTCTGGCATCTCGT *-3’*, 5’- ACCATCCCACTGTCTGTCTG-3’
*vegfc:* 5’-ATGCACTTATTTGGATTTTCTGTCTTCT-3’, 5’-GTCCAGTCTTCCCCAGTATG-3’ *lyve1*: 5’- AGACGTGGGTGAAATCCAAG -3’, 5’- GATGATGTTGCTGCATGTCC-3’
*nr2f2*^47^.

For detection of p53 and p27 embryos were fixed overnight in 4% PFA, washed in methanol, incubated 1 hr. in 3% H2O2 on ice, and stored in methanol at −20C. For p27 immunostaining, embryos were incubated in 150 mM Tris-HCl at pH 9.0 for 5 min, heated at 70C for 15 min and permeabilized in cold acetone at −20C for 20 min. For p53 staining, embryos were digested with 50mg/ul PK followed by 20 min PFA fixation. For both antibodies, embryos were washed in blocking solution containing 0.1% Triton X-100, 0.1% tween, 1% bovine serum albumin and 10% goat serum for 3hrs. at 4C, and incubated with αp53 (55915s, Anaspec, 1:300) or αp27 (sc-528, Santa Cruz, 1:200) antibodies, along with αGFP-biotin (ab6658, Abcam, 1:300) for 3 days at 4C. Samples were then washed with maleic buffer (150Mm maleic acid, 100mM NaCl, 0.1% Tween20, pH 7.4), blocked in maleic buffer containing 2% blocking reagent (Roche, 11-096176-001), and incubated with goat anti-rabbit IgG–horseradish peroxidase (Jackson 1:500) for 2.5 days at 4C for TSA signal amplification, followed by 3 hrs incubation with TSA Plus Cyanine 3 reaction (SAT704A001EA, Perkin Elmer). To recover the endogenous GFP signal, embryos were incubated in 1:300 streptavidin (Jackson, cat # 016-480-084) for 30 min in RT.

### Microscopy and imaging

Live imaging was performed using Zeiss LSM780 or LSM700 upright confocal microscopes (Carl Zeiss, Jena, Germany) equipped with a water immersed X20 NA 1.0 objective lens. Fluorescent proteins were excited sequentially with single-photon lasers (488nm, 561nm).

For time-lapse imaging embryos were held in an imaging chamber containing egg water supplemented with tricaine 0.016% (100 mg/25 ml in fish water) to inhibit movement, and with PTU (0.002%) to prevent pigment development. Embryos held this way, maintained heartbeat and robust circulation throughout the imaging period (up to 50 hours). Z-stacks were acquired at 2.5 μm increments, every 10 min, in 160-170 planes per stack. 2D-, or 3D-reconstructions of image data were prepared using ImageJ (NIH) or Imaris (Bitplane).

### Image processing

Images were processed off-line using ImageJ (NIH) or Imaris (Bitplane). For 3D co-localization analyses in ECs a new colocalization channel was created using the Imaris ‘Colocalization Module’. Co-localization thresholds were set manually. The images shown in this study are single-view, 2D-reconstructions, of collected z-series stacks.

### Statistical Analyses

Comparison of two samples was done using unpaired two-tailed Student’s t-test assuming unequal variance from at least three independent experiments, unless stated otherwise. Statistical significance for three or more samples was calculated via one-way ANOVA followed by post hoc Tukey’s for multiple comparisons.

All data are reported as mean values ± SEM and were analyzed using Prism 5 software (GraphPad Software, Incorporated, La Jolla, CA, USA).

## Acknowledgements

The authors would like to thank Hila Raviv, Lital Shen and Dean Robinson (Weizmann Institute, Israel) for technical assistance, S. Ben-Dor (Weizmann Institute, Israel) for bioinformatic analysis, Gabriella Almog, Roy Hofi, and Anna Tatarin for superb animal care, D. Kimelman (University of Washington) for providing the *dual fucci* transgenic line and *dual fucci* plasmid, T. Look (Dana Farber Cancer Institute, Boston) for providing *p53^m214k^* mutant fish, B. Hogan (University of Melbourne) for providing the *p5E-lyve1* construct. The authors are grateful to all the members of the Yaniv lab for discussion, technical assistance and continuous support. This work was supported in part by European Research Council (818858) to KY, Israel Science Foundation (861/2013) to KY, Binational Science Foundation (2015289) to KY and NDL, Minerva Foundation (712610) to KY, the H&M Kimmel Inst. for Stem Cell Research, the Estate of Emile Mimran (SABRA program). K.Y. is the incumbent of the Enid Barden and Aaron J. Jade Professorial Chair and is supported by the Willner Family Center for Vascular Biology; the estate of Paul Ourieff; the Carolito Stiftung; Lois Rosen, Los Angeles, CA; Edith Frumin; the Fondazione Henry Krenter; the Wallach Hanna & Georges Lustgarten Fund, the Polen Charitable Trust and the Daniel Shapiro Cardiovascular Fund.

## Author Contributions

Conceptualization, A.J.-V. and K.Y.; Methodology, A.J.-V. and G.H.; Investigation, A.J.-V., N.M., D.S., G.H. and M.S.; Writing – Original Draft, A.J.-V.; Writing – Review & Editing, A.J.-V., M.S., N.D.L. and K.Y.; Visualization, A.J.-V. and K.Y.; Supervision, K.Y.; Funding Acquisition, K.Y.

## Notes

### Competing Interest Statement

The authors have declared no competing interest.

