## Supplemental Figures for "VEGFC induced cell cycle arrest mediates sprouting and differentiation of venous and lymphatic endothelial cells"

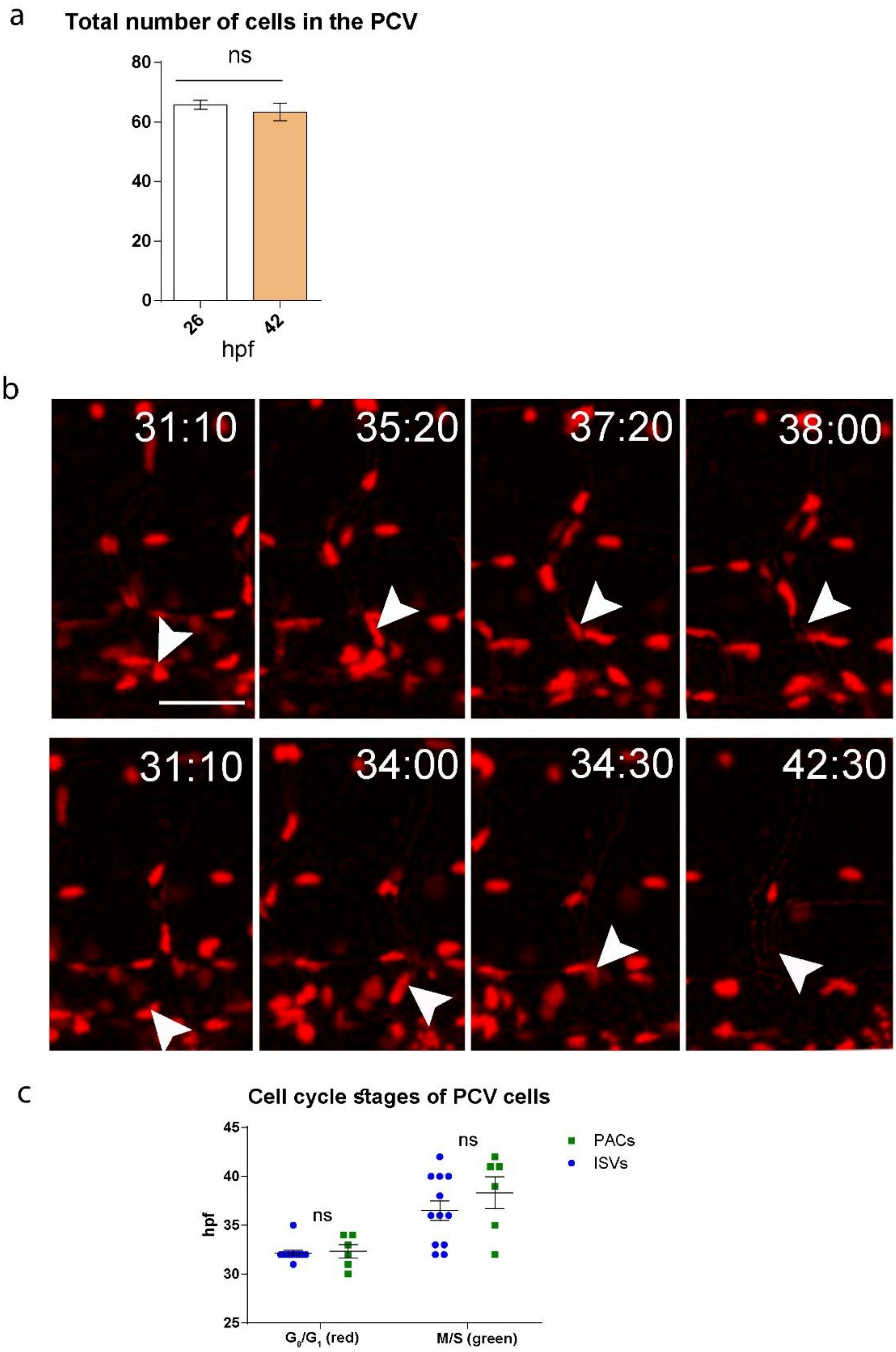


**Figure S1:** (**a**) Total number of ECs within the PCV (10 segments) at 26 hpf and 48 hpf (n=4). (**b**) Red channel of selected confocal snapshots from a time-lapse series of a *Tg(fli1:gal4;uasKaede;uas:fucci)* embryo shown in Figure 1h-o. (**c**) Timing of cell cycle re-entry is vessel type (VEC vs. LEC) independent ($n_{vISV}$=.12, $n_{PAC}$=6). Scale bar=40µm; ns= not significant.

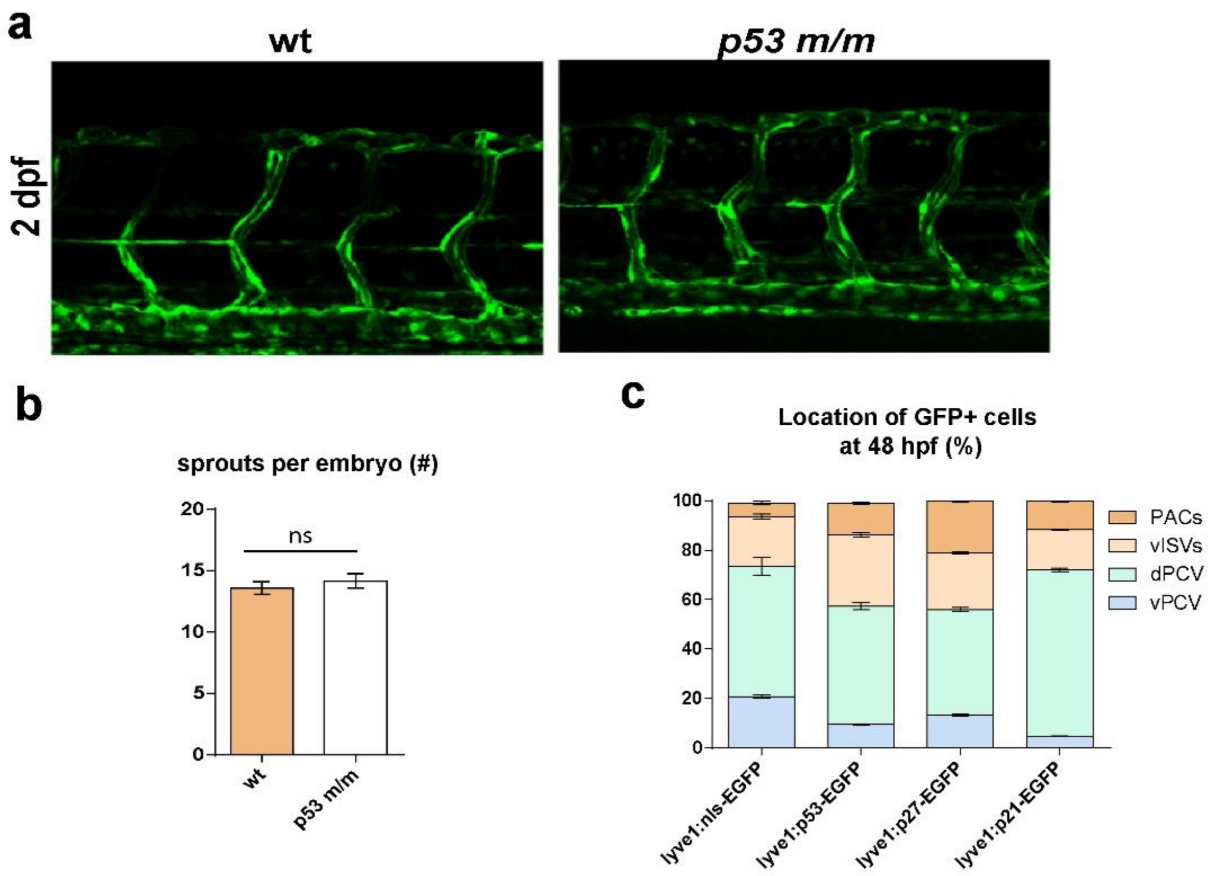


**Figure S2:** (**a**) Confocal images of WT and *p53*$^{m/m}$;*Tg(fli1:EGFP)* embryos at 48 hpf showing normal formation of trunk vasculature in *p53*$^{m/m}$. (**b**) Number of PCV sprouts (vISVs and PACs) in WT vs. *p53*$^{m/m}$ *Tg(fli1:EGFP)* embryos at 48 hpf ($n_{WT}$=12, $n_{p53m/m}$=12). (**c**) Spatial distribution of GFP+ cells in 48 hpf *Tg(fli1:DsRed)* embryos injected with *lyve1:nEGFP* (n=12), *lyve1:p53-EGFP* (n=30), *lyve1:p27-EGFP* (n=12) or stably expressing *lyve1:p21-EGFP*.

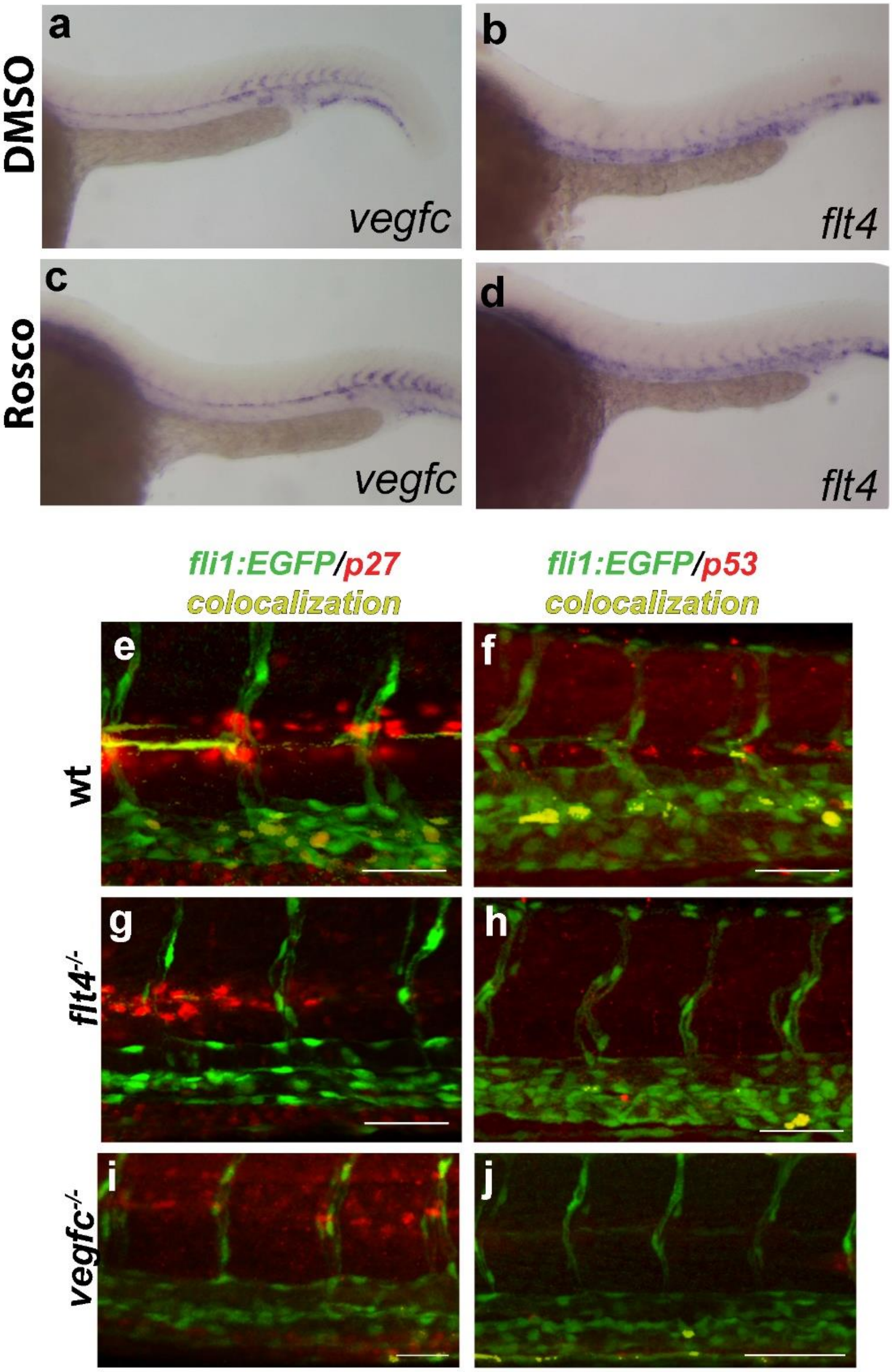


**Figure S3:** (**a**-**d**) *In situ* hybridization on 26 hpf embryos showing no differences in *vegfc* (**a**,**c**) and *flt4* (**b**,**d**) expression following Roscovitine or DMSO treatment.(**e**-j) Confocal images of WT (**e**,**f**), *flt4*$^{-/-}$ (**g**,**h**) and *vegfc*$^{-/-}$ (**i**,**j**) *Tg(fli1:EGFP)*$^{yl}$ embryos at 48 hpf stained with p27 or p53 antibodies (red channels represent immunostaining; co-localization with ECs is highlighted in yellow). Scale bars=70 µm.

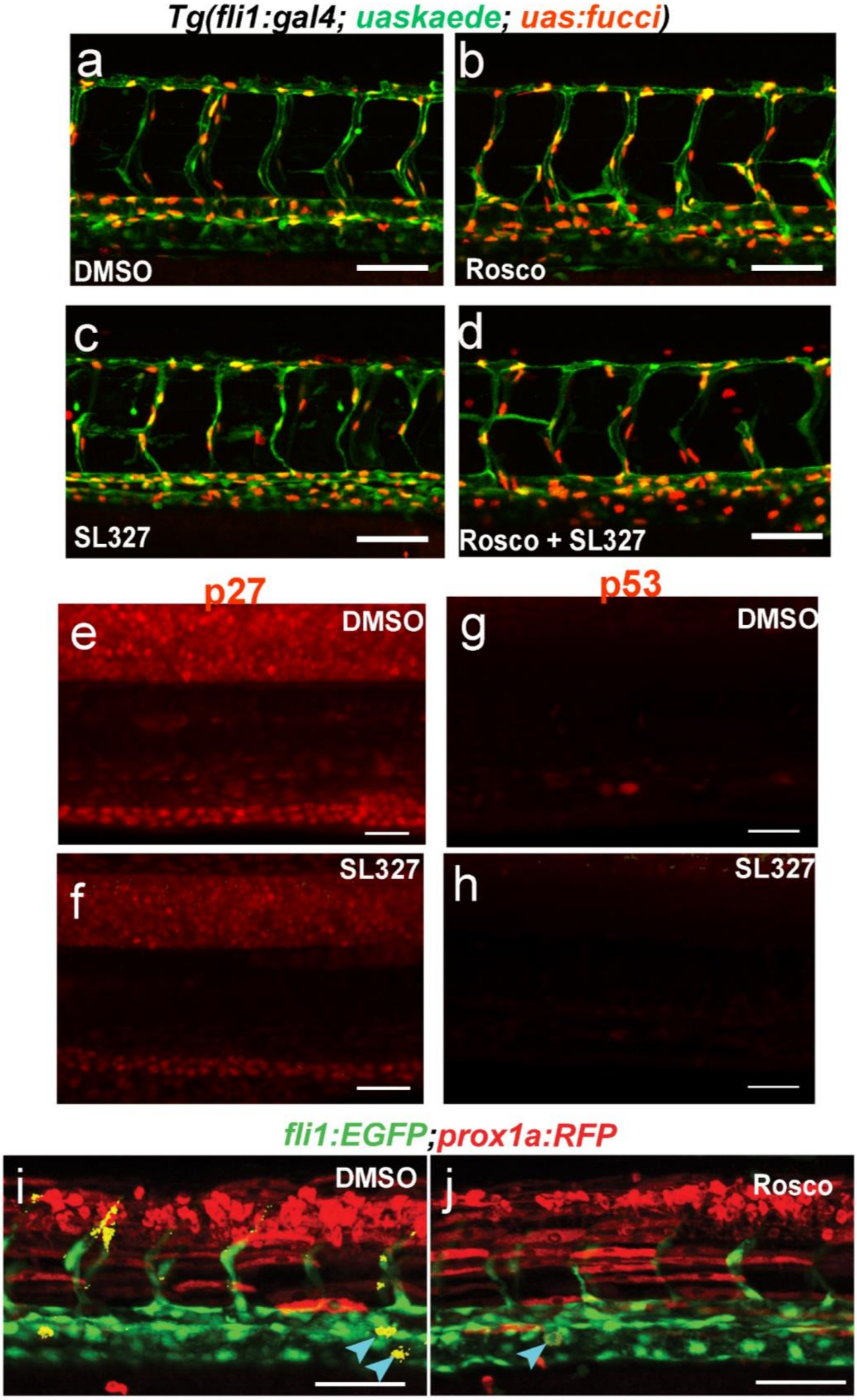


**Figure S4:** (**a-d**) confocal images of 48 hpf *Tg(fli1:gal4;uasKaede;uas:fucci)* embryos treated with DMSO (**a**), Roscovitine (**b**), SL327 (**c**) or Roscovitine+SL327(**d**). (**e-h**) Confocal images (red channel) of *Tg(mrc1a:EGFP)* embryos at 34 hpf, showing reduced expression of p27 (**e, f**) and p53 (**g,h**) following SL327 treatment . (**i,j**) Confocal images of *Tg(fli1:EGFP;prox1a:kalt4:UAS:uncTagRFP)* embryos treated with DMSO (**i**) or Roscovitine (**j**) showing reduction in the number of prox1+ cells (blue arrowheads) within the PCV upon Roscovitine treatment. Co-localization is highlighted in yellow. Scale bars: **a-d, i-j**=70µm, **e-h**=30µm
